# Asynchronous and slow-wave oscillatory states in connectome-based models of mouse, monkey and human cerebral cortex

**DOI:** 10.1101/2023.08.03.551869

**Authors:** Maria Sacha, Jennifer S. Goldman, Lionel Kusch, Alain Destexhe

## Abstract

Thanks to the availability of connectome data that map connectivity between multiple brain areas, it is now possible to build models of whole brain activity. At the same time, advances in mean-field techniques have led to biologically based population models that integrate biophysical features such as membrane conductances or synaptic conductances. In this paper, we show that this approach can lead to brain-wide models of mouse, macaque, and human. We illustrate this approach by showing the transition from wakefulness to sleep simulated with multi-scale models in the three species. We compare the level of synchrony between the three species and found that the mouse brain displays a higher overall synchrony of slow-waves compared to monkey and human brains. We show that these differences are due to the different delays of axonal signal propagation between regions associated to brain-size differences between the species. We also make the program code publicly available, which provides a set of open-source tools for simulating large-scale activity in the cerebral cortex of mouse, monkey, and human.

## 1. Introduction

Brain activity can display widely different states of activity, ranging from the two extremes of asynchronous activity, typical of the aroused brain, and slow and synchronized oscillatory states, reminiscent of slow-wave sleep or anesthesia [1,2]. To model such “macroscopic” states of brain activity, one should ideally take into account “microscopic” features that are key to the genesis of brain states. Therefore, models that can account for biophysical features of brain states must be multi-scale. At the microscopic scale, the genesis of neural activity depends on membrane conductances, such as those responsible for spike-frequency adaptation, which regulate transitions between activity states in cortical slices [3]. The macroscopic features of neural activity between brain states may also crucially depend on recurrent connectivity between excitatory and inhibitory neuron types in cerebral cortex. Here, computational models have been successful to simulate such activity states, either the asynchronous-irregular state [3–6], or the slow-wave state with “Up” and “Down” state dynamics [3,7,8]. Few models could simulate both states, however, because their transition depends on the presence of spike-frequency adaptation conductances. These transitions can be modeled either by Hodgkin-Huxley type models [3] or by integrate-and-fire type models that include spike-frequency adaptation such as the Adaptive Exponential (AdEx) model [8].

Although the genesis of asynchronous and slow oscillatory states can be well modeled at the local network scale, as typically based on *in vitro* activity [9], it is less clear how such activity is organized at larger scales. Here, the activities generated at local scale in each brain area interact through the long-range inter-areal connectivity. The precise pattern of inter-areal connectivity is provided by the connectome, which has been published open-access for a number of species such as mouse [10], macaque monkey [11] and human [12]. So the information is available to construct large-scale models that will include all brain areas and their connectivity. However, simulating such models at the cellular scale represents a huge investment of computational resources, and is out of reach for most researchers having no access to such resources.

An alternative approach is to simulate brain activity using population models, which are much less demanding on computational resources. However, such models need to have enough biological realism to include the relevant biophysical mechanisms necessary to generate brain states, as described above, such as membrane conductances, synaptic receptor types, etc. We will use here an approach that was provided in a recent series of papers [6,8,13,14]. A mean-field model was first derived for AdEx spiking networks [6,8] and successfully generated both asynchronous and synchronized slow-wave dynamics, and their transition by controlling spike-frequency adaptation, as in the experiments. This AdEx mean-field model is much faster to simulate than the spiking networks. Next, such mean-field models were integrated into large-scale models of the human brain [13,14] and could successfully simulate asynchronous states, similar to wakefulness, and slow-wave oscillatory states, similar to slow-wave sleep, as well as their responsiveness to external stimuli (for a similar approach using current-based models, see [15]).

In the present paper, we extend this approach to two species, mouse and macaque monkey. We show the integration of the AdEx mean-field models into large-scale models of the entire cerebral cortex, comparing three species, mouse, monkey and human. In particular, we will illustrate the simulated brain dynamics for asynchronous and slow-wave activity, across the whole cerebral cortex.

## 2. Materials and Methods

We used three types of models, a network of spiking neurons, a mean-field model of this network, and a network of mean-field models implemented in The Virtual Brain (TVB). We describe here these models successively.

### 2.1. Spiking network model

We considered networks of integrate-and-fire neuron models displaying spike-frequency adaptation, based on two previous papers [6,16]. We used the Adaptive Exponential (AdEx) integrate-and-fire model [17]. We considered a population of *N* = 10^4^ neurons randomly connected with a connection probability of *p* = 5%. We considered excitatory and inhibitory neurons, with 20% inhibitory neurons. The AdEx model allows us to define two cell types, “regular-spiking” (RS) excitatory cells, displaying spike-frequency adaptation, and “fast spiking” (FS) inhibitory cells, with no adaptation. The dynamics of these neurons is given by the following equations:

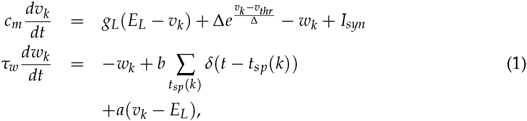

where *c*_*m*_ = 200 pF is the membrane capacitance, *v*_*k*_ is the voltage of neuron *k* and, whenever *v*_*k*_ > *v*_*thr*_ = −50 mV at time *t*_*sp*_(*k*), *v*_*k*_ is reset to the resting voltage *v*_*rest*_ = −65 mV and fixed to that value for a refractory period *T*_*refr*_ = 5 ms. The leak term *g*_*L*_ had a fixed conductance of *g*_*L*_ = 10 nS and the leakage reversal *E*_*L*_ was of −65 mV. The exponential term had a different strength for RS and FS cells, i.e. Δ = 2mV (Δ = 0.5mV) for excitatory (inhibitory) cells. Inhibitory neurons were modeled as fast spiking FS neurons without adaptation (*a* = *b* = 0 for all inhibitory neurons) whereas excitatory regular spiking RS neurons had a lower level of excitability due to the presence of adaptation (while *b* varied in our simulations we fixed *a* = 4 nS and *τ*_*w*_ = 500 ms unless otherwise specified). The synaptic current *I*_*syn*_ received by neuron *i* is the result of the spiking activity of all neurons *j* ∈ pre(*i*) pre-synaptic to neuron *i*. This current can be decomposed in the synaptic conductances evoked by excitatory E and inhibitory I pre-synaptic spikes

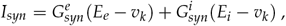

where *E*_*e*_ = 0mV (*E*_*i*_ = −80mV) is the excitatory (inhibitory) reversal potential. Excitatory synaptic conductances were modeled by a decaying exponential function that sharply increases by a fixed amount *Q*_*E*_ at each pre-synaptic spike, i.e.:

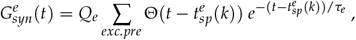

where Θ is the Heaviside function, *τ*_*e*_ = *τ*_*i*_ = 5ms is the characteristic decay time of excitatory and inhibitory synaptic conductances, and *Q*_*e*_ = 1 nS (*Q*_*i*_ = 5 nS) the excitatory (inhibitory) quantal conductance. Inhibitory synaptic conductances are modeled using the same equation with *e* → *i*. This network displays two different states according to the level of adaptation, *b* = 5 pA for asynchronous-irregular states, and *b* = 60 pA for Up-Down states (Fig. 1A; see [6] for details).

**Figure 1.**
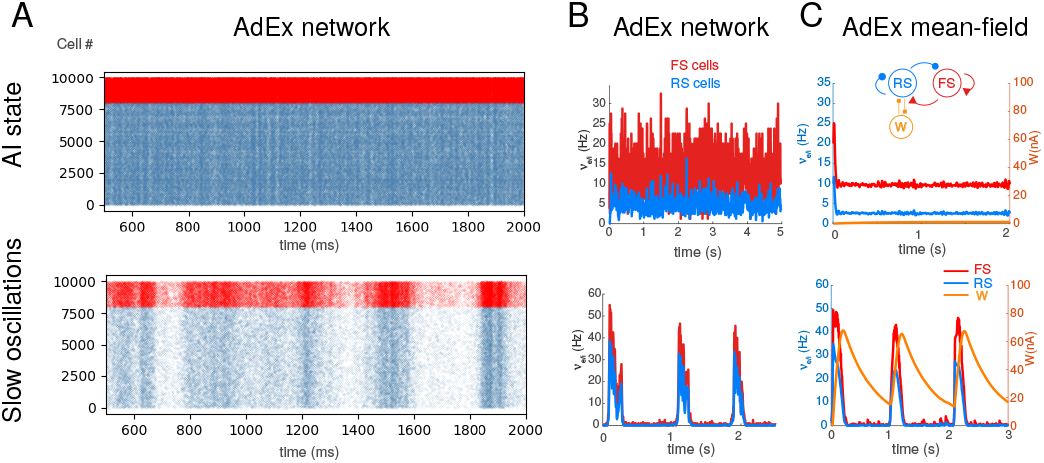
Asynchronous-irregular and slow oscillatory states in AdEx networks and mean-field models. A. Raster plots of excitatory RS (blue) and inhibitory FS (red) AdEx neurons in a network during asynchronous-irregular (AI) state (top), and slow oscillations with Up and Down states (bottom). The two states differed from the values of adaptation parameter *b* (*b*=1 pA for AI state, and 20 pA for slow oscillations shown here). B. Corresponding mean firing rates of the two populations. C. Mean-field model of AdEx networks with adaptation (scheme in inset). The mean-field reproduces the two states and the corresponding firing rate variations. Modified from [8,13,18].

### 2.2. Mean-field models

We considered a population model of a network of AdEx neurons, using a Master Equation formalism originally developed for balanced networks of integrate-and-fire neurons [19]. This model was adapted to AdEx networks of RS and FS neurons [6], and later modified to include adaptation [8]. The latter version is used here, which corresponds to the following equations:

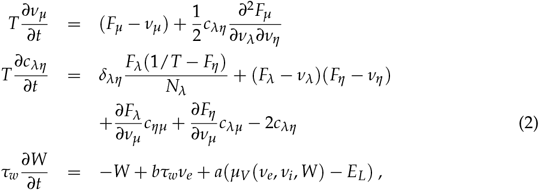

where *μ* = {*e, i*} is the population index (excitatory or inhibitory), *ν*_*μ*_ the population firing rate and *c*_*λη*_ the covariance between populations *λ* and *η. W* is a population adaptation variable [8]. *δ*_*λη*_=1 if *λ* = *η* and zero otherwise. The function *F*_*μ*= {*e,i*}_ = *F*_*μ*= {*e,i*}_ (*ν*_*e*_, *ν*_*i*_, *W*) is the transfer function which describes the firing rate of population *μ* as a function of excitatory and inhibitory inputs (with rates *ν*_*e*_ and *ν*_*i*_) and adaptation level *W*. These functions were estimated previously for RS and FS cells and in the presence of adaptation [8].

At the first order, i.e. neglecting the dynamics of the covariance terms *c*_*λη*_, this model can be written simply as:

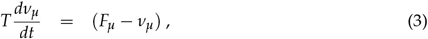

together with Eq. 3. This system is equivalent to the well-known Wilson-Cowan model [20], with the specificity that the functions *F* need to be obtained according to the specific single neuron model under consideration. These functions were obtained previously for AdEx models of RS and FS cells [6,8] and the same are used here.

For a cortical volume modeled as a two populations of excitatory and inhibitory neurons, the equations (at first order) can be written as:

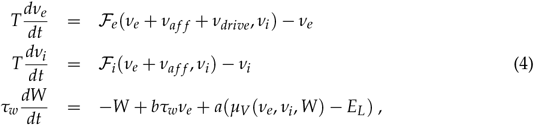

where *ν*_*aff*_ is the afferent thalamic input to the population of excitatory and inhibitory neurons and *ν*_*drive*_ is an external noisy drive. The function *μ*_*V*_ is the average membrane potential of the population and is given by

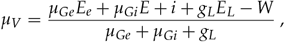

where the mean excitatory conductance is *μ*_*Ge*_ = *ν*_*e*_*K*_*e*_*τ*_*e*_*Q*_*e*_ and similarly for inhibition.

This system describes the population dynamics of a single isolated cortical column, and was shown to closely match the dynamics of the spiking network (Fig. 1; [8]).

### 2.3. Networks of mean-field models

Extending our previous work at the mesoscale [8,21] to model large brain regions, we define networks of mean-field models, representing interconnected cortical columns (each described by a mean-field model). For simplicity, we considered only excitatory interactions between cortical columns, while inhibitory connections remain local to each column. The equations of such a network, expanding the two-population mean-field (Eq. 4), are given by:

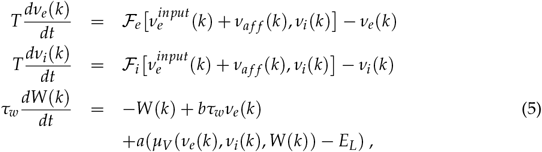

where *ν*_*e*_(*k*) and *ν*_*i*_(*k*) are the excitatory and inhibitory population firing rates at site *k*, respectively, *W*(*k*) the level of adapation of the population, and 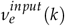 is the excitatory synaptic input. The latter is given by:

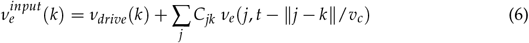

where the sum runs over all nodes *j* sending excitatory connections to node *k*, and *C*_*jk*_ is the strength of the connection from *j* to *k* (and is equal to 1 for *j* = *k*). Note that *ν*_*e*_(*j, t* − ∥*j* − *k*∥/*v*_*c*_) is the activity of the excitatory population at node *k* at time *t* − ∥*j* − *k*∥/*v*_*c*_ to account for the delay of axonal propagation. Here, ∥*j* − *k*∥ is the distance between nodes *j* and *k* and *v*_*c*_ is the axonal propagation speed.

As detailed in a previous study [14], the spike-frequency adaptation parameter in the model can be linked to the neuromodulatory drive. From a biological perspective, during wakefulness, increased concentrations of neuromodulators, such as acetylcholine, NE and 5HT, diminish spike-frequency adaptation by down-regulating various K+ channels, which results in a prolonged depolarization of the neurons and enables the emergence of asynchronous and irregular firing [22]. On the contrary, during unconsciousness, the lower concentrations of neuromodulators leave the K+ channels open. The adaptation build-up and its subsequent decay results in synchronous hyperpolarization and depolarization of the neuron’s membrane potential, which in turn leads to the generation of slow-wave dynamics. *b* controls the strength of spike-frequency adaptation so that augmenting *b* corresponds to a reduced neuromodulatory drive, and switches the activity from asynchronous to slow-waves (see details in [8,13,14]).

### 2.4. Connectomes for the three species

The mouse connectome used here is a parcellation comprising 98 regions. The connectivity matrix was created through the Allen Connectivity Builder which uses high-resolution anterogade tract-tracing data provided by the Allen Institute of Brain Science. The experiments concern source regions only in the right hemisphere, therefore the left hemisphere is build as the mirror image of the right (data are available in: https://zenodo.org/records/8331301, for more information, see ref [23]).

The macaque structural connectivity matrix, consists of 82 nodes and it was generated through a synthesis of axonal tract-tracing and diffusion-weighted imaging data, resulting in a directed and weighted whole-cortex macaque connectome (data available in: https://zenodo.org/records/7011292; for more information, see ref. [24]).

The human connectome includes 68 nodes for which the connection was based on human tractography methods from the Berlin empirical data processing pipeline [25]. Diffusion weighted imaging does not provide information on fiber tracts directionality, but these information can be derived from tracer-studies on macaques, and subsequently mapped on the human brain (data available in https://zenodo.org/records/7574266).

### 2.5. Integration in The Virtual Brain

The integration of networks of mean-field models was done for each species using The Virtual Brain (TVB) simulator (https://www.thevirtualbrain.org/tvb). For the mouse brain, we used “The Virtual Mouse Brain” [23], the monkey brain model was given by the macaque TVB [24], and the TVB model of the human brain was from [26]. These publications should be consulted for details about the connectivity used. The particular implementation of the human TVB was that given in [14], where more details can be obtained.

### 2.6. Analysis

To quantify the amount of synchrony in the TVB model, we computed the Phase-Lag Index (PLI) for each pair of nodes, averaged over the simulation time. The Hilbert transform is first computed to extract the phase *ψ*(*t*) of the time series. From there, the PLI is given by

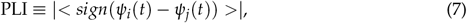

where <·> denotes averaging over time, for nodes *i* and *j* [27]. Note that the PLI takes values between 0 (random phase relations or perfect synchrony) and 1 (perfect phase locking). In this work we report the mean PLI over all time epochs for excitatory and inhibitory firing rates of each region pair for each adaptation value.

## 3. Results

We start by showing the behavior of the mean-field model, then we show its integration in brain-wide models of mouse, monkey and human cerebral cortex.

Figure 1 shows the spiking network model of AdEx neurons, comprising RS (blue) and FS (red) cells, as detailed in previous studies [6,8]. This network can generate asynchronous irregular (AI) activity states (Fig. 1A, top), as well as slow synchronized activity under the form of Up and Down states (Fig. 1A, bottom). The transition from these two states is obtained by strengthening or weakening of the spike-frequency adaptation in RS cells (parameter *b* in the model). The mean rates of activity of these two states are shown in Fig. 1B, respectively. The mean-field model of this AdEx network was derived in two previous publications [6,8] and is shown for these two states in Fig. 1C.

As shown previously [13,14], the AdEx mean-field model was implemented in The Virtual Brain (TVB) platform. This platform consists of a python-based simulation environment [26] which allows the user to create a network of mean-field models (or more generally neural mass models), constrained by connectivity data extracted from a given connectome. The TVB environment can also generate a number of neural signals and be linked to neuro-imaging [29]. Integrating the AdEx mean-field in TVB, leading to the “TVB-AdEx” model [13], has a double advantage. First, the mean-field model is biologically informed, so it has physically-interpretable parameters (such as conductances, synaptic receptor kinetics, etc) that can be changed and directly compared to experiments. Second, the AdEx mean-field can generate states of activity which can be asynchronous or synchronized (see Fig. 1), which is not the case with other neural mass models. This behavior is remarkably robust within a large parameter space of the model, as investigated in detail previously [28] and illustrated in Fig. 2.

**Figure 2.**
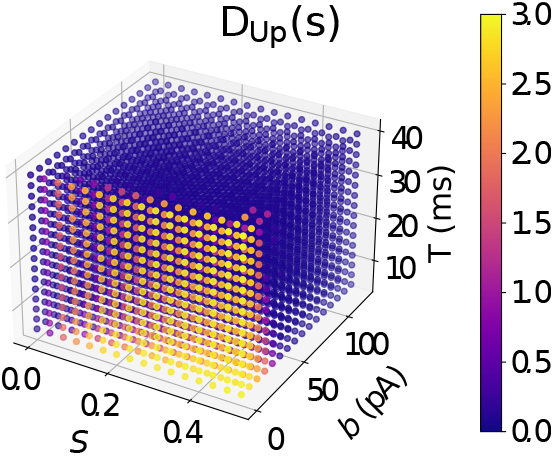
Robustness of asynchronous and synchronized slow oscillatory states in the human TVB-AdEx model. The 3D graph represents the TVB model behavior in a space spanned by three parameters, the connection strength *S*, the level of adaptation *b* and the time scale *T*. The color indicates the duration of the up-states *D*_*up*_ (large durations indicate that the dynamics is sustained and asynchronous). Modified from [28].

The behavior of the TVB-AdEx model is illustrated in Fig. 3 for mouse, monkey and human brains. In all three species, the asynchronous activity of the mean-field model leads to a global dynamics which is also asynchronous (Fig. 3, middle panels). While there were some variations of the absolute amplitudes of the firing rates, the three models displayed consistent asynchronous dynamics. For higher levels of adaptation, the three models displayed slow-wave activity, as in the mean-field model. However, in this case the individual nodes synchronized their slow oscillation, which generated a globally synchronized slow-wave activity across large brain regions. In the case of the human brain, a grid scan of parameters showed that the level of synchrony depends on the parameters, and in particular on the strength of the long-range connectivity [14]. Similar behavior was observed in the three models illustrated here (not shown).

**Figure 3.**
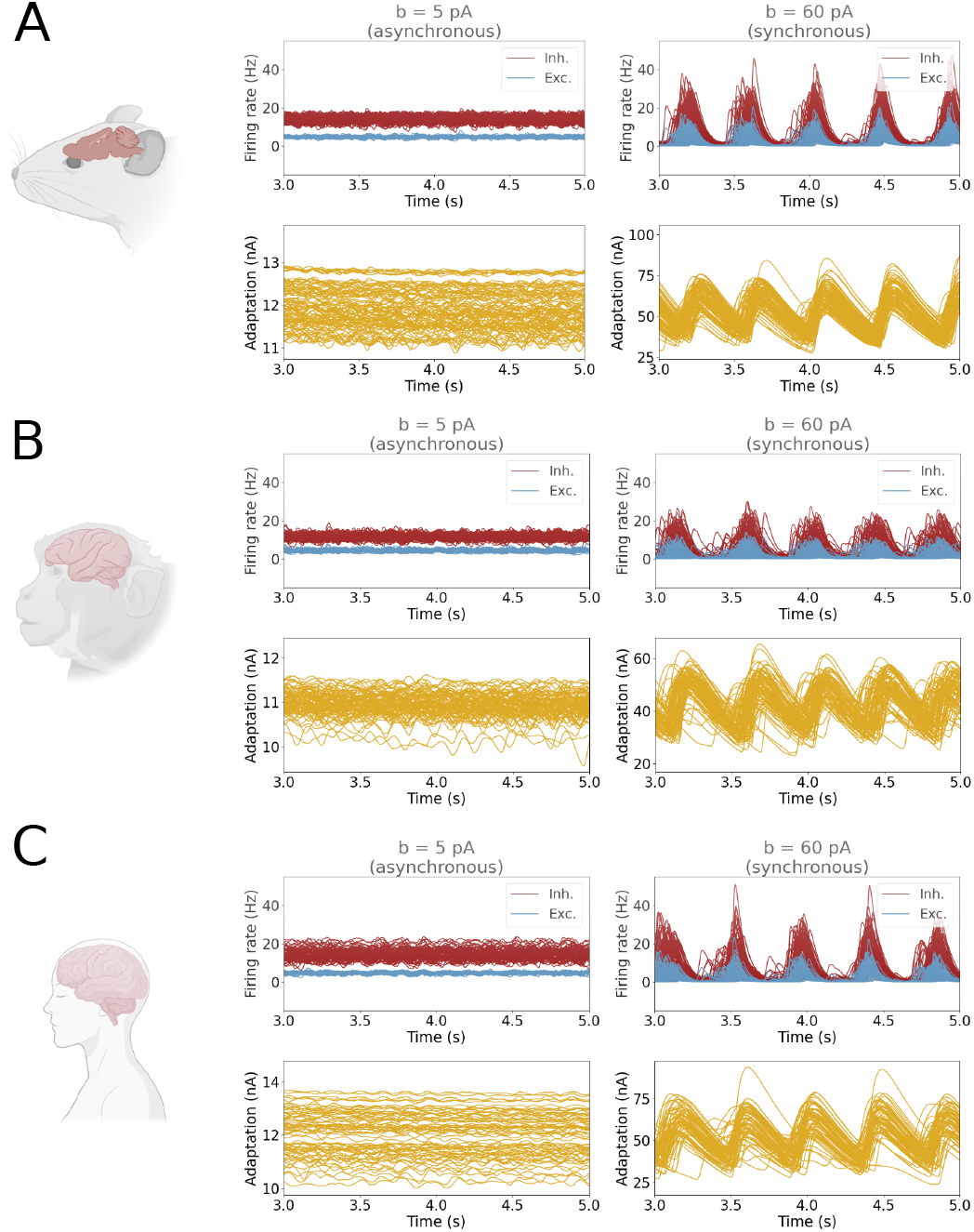
Emergent asynchronous and synchronized slow-wave dynamics in mouse, macaque and human brain models. A: mouse brain, B: macaque brain, C: human brain. In each case, the middle panels show the asynchronous activity resulting from low levels of spike-frequency adaptation (*b*=5 pA), while the right panels show the slow-wave dynamics obtained for higher levels of spike-frequency adaptation (*b*=60 pA). The color code is blue for excitatory firing rates, red for inhibitory firing rates, and yellow for mean adaptation. Panel C was modified from [14].

The comparison of the two states across the models of the three species is further shown in Fig. 4. The transition from asynchronous to slow-wave dynamics occurs at about the same value of adaptation parameter *b* (Fig. 4A). Interestingly, the duration of the down states is predicted to be similar between species. One can also see that the down-state duration increases with *b*, and thus the slow-wave frequency will also decrease proportionally to the level of adaptation. The level of synchrony of the activities is shown in Figs. 4B-C, as measured by the phase-lag index (PLI), for excitatory and inhibitory population activities, respectively. The synchrony increases for excitatory cells as a function of the adaptation parameter *b* (Figs. 4B),. Interestingly, one can see that the level of synchrony is systematically higher for the mouse brain, intermediate for the human brain, and lower for the monkey brain. Because the mouse brain is considerably smaller, the axonal delays are also smaller than the other two models. To test its effect on synchrony, we have matched the delay distributions of the three species. The axonal propagation speed of the mouse and macaque was decreased, resulting in increased delays in these two species that they are similar to the delay distribution in human. In this case, as shown in Fig. 4C, the level of synchrony of the mouse brain diminishes and roughly matches that of the human brain. However, the macaque brain still displays a lower level of synchrony, which may be due to a sparser connectivity. This prediction of the model should be tested by appropriate experiments.

**Figure 4.**
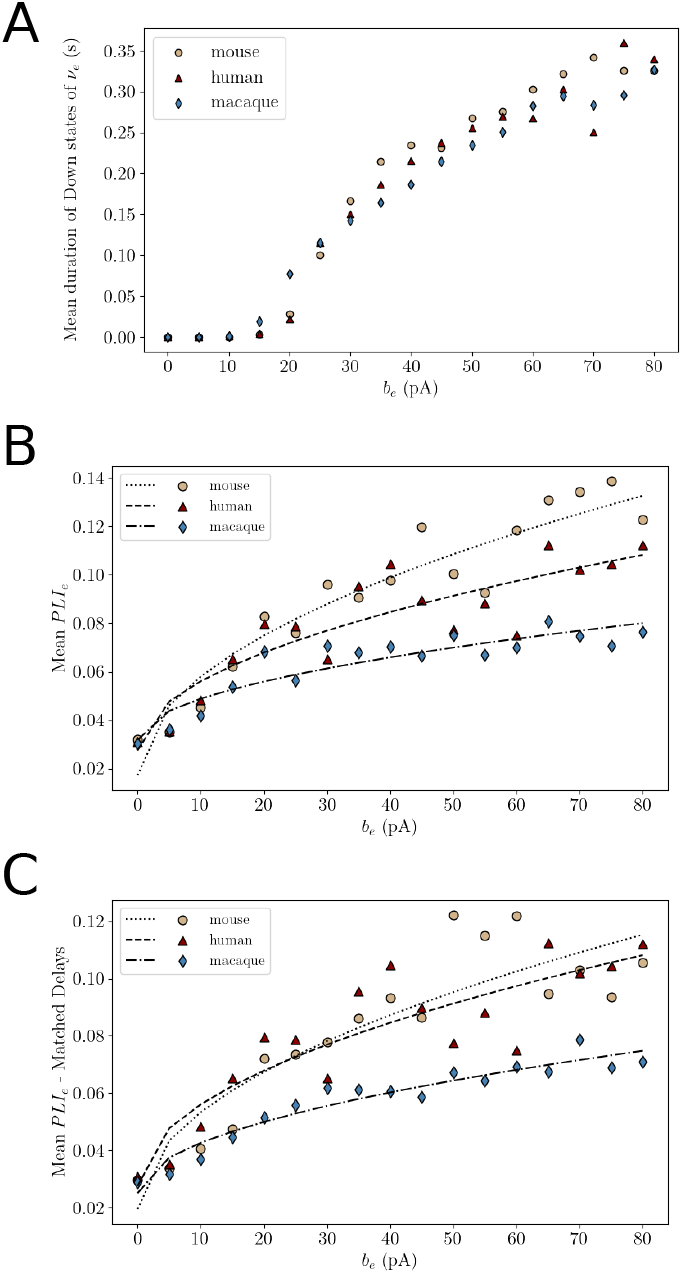
Comparison of the asynchronous and synchronized slow-wave dynamics in mouse, macaque and human brain models. A: Transition between asynchronous and slow-wave dynamics measured by the mean duration of down states. B. Mean phase-lag index (PLI) for excitatory neuron activities for the three species. C. Same as B, but the connection delays were matched between the three connectomes.

## 4. Discussion

In this paper, we have shown the integration of AdEx mean-field models into whole-cortex simulations for three species, mouse, monkey and human. The chosen model is a second-order mean-field model, grounded on biophysically plausible spiking models, and built using a bottom-up approach. The model is able to reproduce asynchronous and slow oscillatory states of neural networks, and therefore permits us to evaluate the emergence of synchronized slow-waves at the whole-brain level across three distinct species. For each species, the simulations exhibit the main features of the TVB-AdEx model, as found earlier [13,14]. In particular, the three models exhibit the same emerging property found in the human brain simulations, namely that when the individual AdEx nodes are set in the asynchronous mode, the ensemble of nodes remains asynchronous. This mode of activity is consistent with the “desynchronized” activity classically found in the awake human brain [1,2] and which is also seen in the human and monkey brain in microelectrode recordings which also display asynchronous-irregular states [30]. Similarly, when the AdEx nodes are set to the Up/Down state mode, the activity over the whole cortex synchronizes into a slow oscillatory mode. Here also, this slow oscillatory mode corresponds to the Up & Down states seen in single units in microelectrode recordings in human and monkey during slow-wave sleep [30,31]. Microelectrode recordings in mice also display the same two states, asynchronous-irregular dynamics during wakefulness and Up & Down state dynamics during slow-wave sleep [32], and similarly in cats [33].

We believe that the simulations explored here, and for which we provide the python code [34], constitute a very useful tool for the community of computational neuroscientists interested in simulating whole-brain dynamics in mouse, monkey and human. We must note here that the code we provide are codes running within the TVB environment [26]. In addition, we also have implemented versions of these models which can be run online, on the EBRAINS platform. This will allow users to run the models and change the parameters, without even installing TVB, which could be a useful tool as well.

Finally, we would like to mention a few possible limitations and extensions to the models explored here. The connectomes used in our study have been derived from different experimental methods (axonal tract-tracing, diffusion weighted images, or a combination of both) and they may exhibit discrepancies in terms of spatial resolution, sensitivity, and accuracy due to inherent differences in the underlying data acquisition and processing. As a direct implication, the connectomes lack consistent directionality (since diffusion-weighted imaging do not provide this information), which may have an effect on the connectivity patterns and their emergent dynamics. It is also important to keep in mind that these are purely cortical models, ignoring the contribution of subcortical structures in the generation of slow-wave activity, with rather coarse brain parcellations that include less than a hundred regions for each species.

A first extension is thus to complete the present model, which is only cortical, by other regions, such as thalamus, hippocampus, basal ganglia, cerebellum, etc. Integrating these regions will require the design of mean-field models specific to each brain region, which was done recently for cerebellum [35] and thalanus [36], and is in progress for other brain regions. The models would also benefit from the use of high resolution parcellations that take into consideration local circuits and dynamics. A second extension is to study other states and other transitions than the ones explored here. For example, there could be oscillatory states (such as beta, or gamma) occurring as responses to sensory stimulation, slow-wave states occurring under anesthesia, or pathological states such as coma or minimal consciousness. To model such states, the procedure is the same as outlined here: start from a spiking network model displaying these new states, then design a mean-field model that can capture this effect, and finally integrate the mean-field at large scales in TVB. We believe that these are interesting directions to explore and that the models proposed here will provide useful tools for this exploration.

## Author Contributions

Conceptualization, J.S.G and A.D; methodology, J.S.G, M.S, L.K.; analysis and preparation of the figures, M.S.; writing of the initial manuscript, A.D.; supervision, project administration and funding acquisition, A.D. All authors have discussed the results and agreed to the published version of the manuscript.

## Data Availability Statement

The source code of all simulations shown in this article is available online. A repository is available in *Zenodo* [34], and the code can be run on-line using the simulation capabilities offered by the Human Brain Projectâs EBRAINS neuroscience research infrastructure (https://ebrains.eu and https://ebrains.eu/service/the-virtual-brain).

## Acknowledgments

We thank Bahar Hazal Yalçinkaya, Trang-Anh Nghiem, David Aquilue, Kevin Ancourt and Viktor Jirsa for their help, support and discussion.

## Disclaimer/Publisher’s Note

The statements, opinions and data contained in all publications are solely those of the individual author(s) and contributor(s) and not of MDPI and/or the editor(s). MDPI and/or the editor(s) disclaim responsibility for any injury to people or property resulting from any ideas, methods, instructions or products referred to in the content.

